# Vesicular stomatitis virus chimeras expressing the Oropouche virus glycoproteins elicit protective immune responses in mice

**DOI:** 10.1101/2021.02.19.432025

**Authors:** Sarah Hulsey Stubbs, Marjorie Cornejo Pontelli, Nischay Mishra, Changhong Zhou, Juliano de Paula Souza, Rosa Maria Mendes Viana, W. Ian Lipkin, David M. Knipe, Eurico Arruda, Sean P. J. Whelan

## Abstract

Oropouche virus (OROV) infection of humans is associated with a debilitating febrile illness that can progress to meningitis or encephalitis. First isolated from a forest worker in Trinidad and Tobago in 1955, the arbovirus OROV has since been detected throughout the Amazon basin with an estimated 500,000 human infections. Like other members of the family *Peribunyaviridae*, the viral genome exists as 3 single-stranded negative-sense RNA segments. The medium sized segment encodes a viral glycoprotein complex (GPC) that is proteolytically processed into two viral envelope proteins Gn and Gc responsible for attachment and membrane fusion. There are no therapeutics or vaccines to combat OROV infection, and we have little understanding of protective immunity to infection. Here we generated a replication competent chimeric vesicular stomatitis virus (VSV), in which the endogenous glycoprotein was replaced by the GPC of OROV. Serum from mice immunized with VSV-OROV specifically neutralized wild type OROV, and using peptide arrays we mapped multiple epitopes within an N-terminal variable region of Gc recognized by the immune sera. VSV-OROV lacking this variable region of Gc was also immunogenic in mice producing neutralizing sera that recognize additional regions of Gc. Challenge of both sets of immunized mice with wild type OROV shows that the VSV-OROV chimeras reduce wild type viral infection and suggest that antibodies that recognize the variable N-terminus of Gc afford less protection than those that target more conserved regions of Gc.

**Importance:** Oropouche virus (OROV), an orthobunyavirus found in Central and South America, is an emerging public health challenge that causes debilitating febrile illness. OROV is transmitted by arthropods, and increasing mobilization has the potential to significantly increase the spread of OROV globally. Despite this, no therapeutics or vaccines have been developed to combat infection. Using vesicular stomatitis (VSV) as a backbone, we developed a chimeric virus bearing the OROV glycoproteins (VSV-OROV) and tested its ability to elicit a neutralizing antibody response. Our results demonstrate that VSV-OROV produces a strong neutralizing antibody response that is at least partially targeted to the N-terminal region of Gc. Importantly, vaccination with VSV-OROV reduces viral loads in mice challenged with wildtype virus. This data provides the first evidence that targeting the OROV glycoproteins may be an effective vaccination strategy to combat OROV infection.

## Introduction

Oropouche virus (OROV), first isolated in 1959 from a forest worker in Trinidad and Tobago, causes a debilitating febrile illness in humans that can progress to meningitis or encephalitis (1, 2). OROV is the most prevalent arbovirus after dengue in Brazil (3, 4), and is currently circulating in Argentina, Bolivia, Colombia, Ecuador, and Venezuela (5–7). In urban areas across the Amazon region seroprevalence rates of up to 15-33%, suggest that OROV infection is underappreciated (8, 9). The virus infects a broad range of species and has both a sylvatic and urban cycle. During the sylvatic cycle, the virus infects sloths, monkeys, rodents, and birds, with *Coquillettidia venezuelensis* and *Ochlerotatus serratus* serving as vectors (1). In the urban cycle, *Culicoides paraensis* and *Culex quinquefasciatus* serve as vectors of OROV (1), with human infections paralleling increases in the vector population during the rainy season (10). Infection is predominantly seen in individuals returning from forested areas (11), with limited evidence of direct human to human transmission (12). Climate change, expansion and dissemination of vectors, globalization of travel, and habitat loss likely contribute to increases in OROV infections (13). Despite the growing dissemination of OROV, and the risks posed by this emergent threat, there are currently no therapeutics to combat OROV infection.

Oropouche virus, a member of the family *Peribunyaviridae*, contains a single-stranded, negative-sense RNA genome divided on three segments providing a total genome size of 11, 985 nucleotides. The large segment (l) encodes the large protein (L) which acts as the RNA dependent RNA polymerase, the medium segment (m) encodes the viral glycoprotein complex (GPC), and the small segment (s) encodes the nucleocapsid protein that sheaths the genomic and antigenomic RNA. Two non-structural proteins, NSs and NSm are coded by the small and medium segments, respectively. The GPC is synthesized as a single polyprotein that undergoes co-translational cleavage by host-cell proteases into the two glycoproteins, Gn and Gc and liberates the intervening NSm protein (14). The Gc protein acts as the viral fusogen (15), and Gn mediates attachment and shields and protects Gc from premature triggering (16). Extrapolating from studies with La Crosse (17–20) and Schamllenberg (16, 21) viruses, both in the same orthobunyavirus genus of the *Peribunyaviridae*, antibodies against the glycoproteins may protect against infection and are therefore an important objective of vaccine strategies against OROV.

Vesicular stomatitis virus (VSV), the prototype of the family *Rhabdoviridae*, infects cells by a single attachment and fusion glycoprotein (G) (22). The development of reverse genetic approaches to manipulate the VSV genome (23) permitted replacement of the attachment and fusion machinery with those of heterologous lipid enveloped viruses (24–28). In the case of Zaire ebolavirus, the resulting VSV-ZEBOV chimera is a live attenuated vaccine, ERVEBO^®^ that is approved for use in humans (29–31). Analogous vaccine candidates are in pre-clinical and clinical development for multiple enveloped viruses (25, 27, 32), including severe acute respiratory syndrome 2 coronavirus (33). In addition to providing a platform for development of candidate vaccines, the VSV-chimeric viruses have also permitted investigation of viral tropism, cellular entry pathways, and helped understand correlates of immune protection (30, 34–38).

Building on this proven approach, we generate a VSV-chimera in which the native glycoprotein gene is replaced by the GPC of OROV. The resulting VSV-OROV chimera replicates to high-titers in BSRT7 cells in culture, and efficiently incorporates the Gn and Gc of OROV into particles. Following single dose or prime-boost intramuscular injection of mice, VSV-OROV elicits the production of immune specific sera that neutralize wild-type OROV. Using peptide arrays corresponding to the OROV glycoproteins we find significant reactivity to the more variable N-terminal domain of Gc that precedes the domains that are structurally homologous to other class II fusion proteins. We generate two additional VSV-OROV chimeras lacking portions of the variable N-terminus of Gc, termed the “head” and “stalk” domain. Although both variants yielded replication competent virus deletion of the stalk domain led to the accumulation of mutations in Gc. Immunization of mice with the VSV-OROV lacking the head domain of Gc generated immune sera reactive with new regions of Gc. Challenge studies demonstrate that immunization with VSV-OROV or the chimera lacking the head domain of Gc offer protection against wild type OROV infection as evidenced by weight loss, temperature, and viral burden. This study demonstrates that VSV-OROV immunization induces neutralizing serum responses that can protect mice against challenge with wild type OROV.

## Results

### Construction and characterization of VSV-Oropouche chimeras

Orthobunyavirus virions incorporate two viral glycoproteins on their surface, Gn and Gc that mediate binding and entry into the cell. Gn and Gc are synthesized as part of a polyprotein precursor GPC. We engineered an infectious molecular clone of VSV that expresses eGFP as a marker of infection (39), to replace the native attachment and fusion glycoprotein with the GPC polyprotein of OROV (Figure 1A). Using established procedures, we recovered a chimeric virus, VSV-OROV, capable of autonomous replication as evidenced by plaque formation (Figure 1A). Kinetic analysis of the yields of infectious VSV-OROV identify a 1 log reduction in viral growth compared to VSV-eGFP (Figure 1A). Analysis of the protein composition of purified virions by SDS-PAGE demonstrates that Gn and Gc are incorporated into VSV-OROV in place of VSV G (Figure 1B). VSV-OROV particles retain the classic bullet shape of VSV as evidenced by negative-stain electron microscopy (Figure 1C) and are visually decorated with spikes consistent with incorporation of the OROV glycoproteins (Figure 1C). As insertion of the M ORF of OROV increases the genome length of VSV by approximately 2500 nucleotides, the VSV-OROV particles are approximately 34 nm longer than VSV particles (Figure 1D).

**Figure 1:**
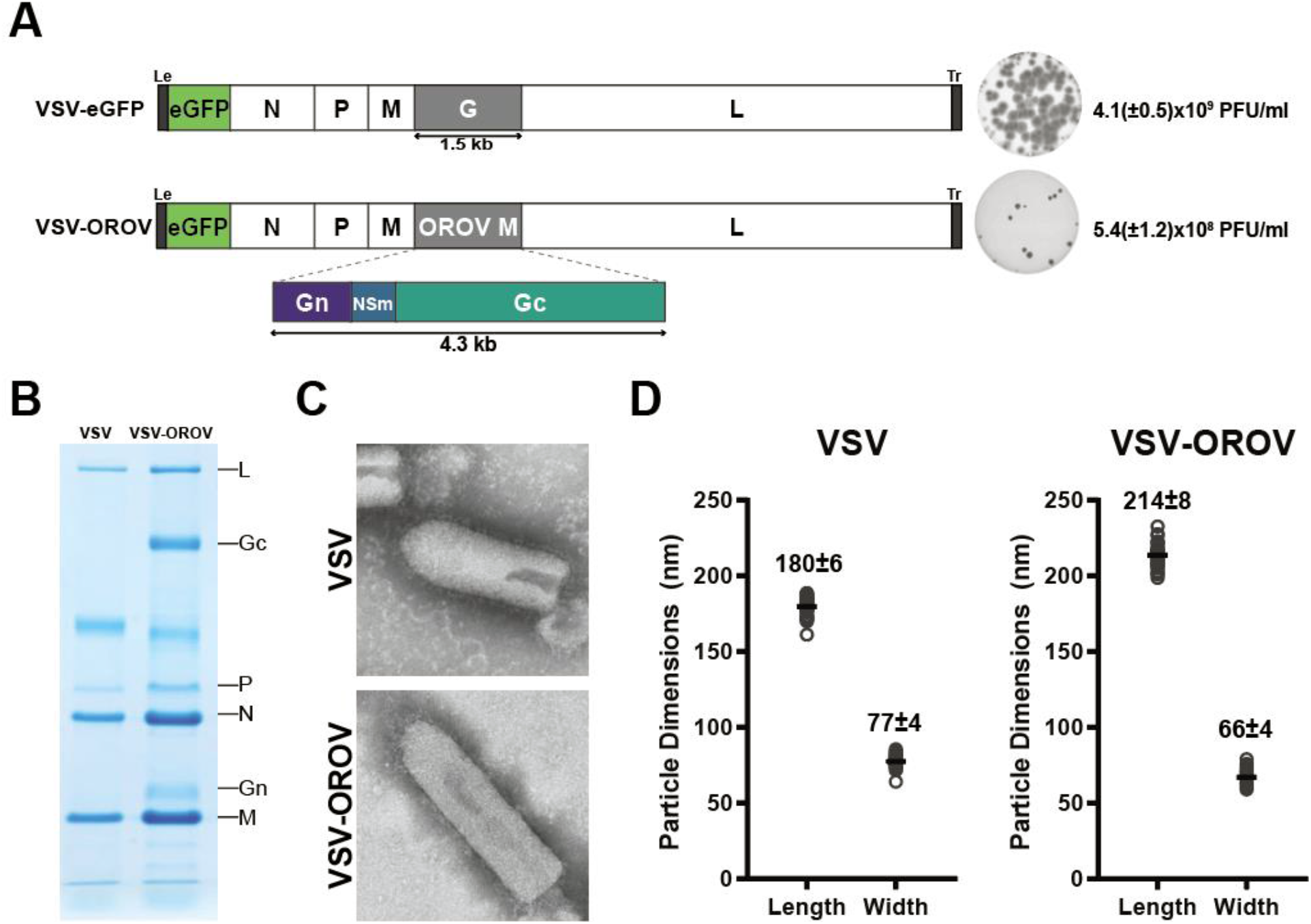
Generation and characterization of VSV-OROV. (A) Genomic organization of VSV-eGFP and VSV-OROV. Viral genomes are shown in the 3’ to 5’ orientation. The five viral genes, N (nucleocapsid), P (phosphoprotein), M (matrix), G (glycoprotein), and L (large polymerase), are shown. Enhanced green fluorescent protein (eGFP) is located in the first position and serves as a marker for infection. The VSV glycoprotein was replaced with the gene encoding the OROV M segment to generate VSV-OROV. Representative plaque assays are shown on the right as well as the end point titers in BSRT7 cells. (B) SDS-PAGE analysis of purified virions stained with Coomassie blue. Viral proteins are indicated on the right. (C) Electron micrographs of purified virions stained with 2% PTA. (D) Measurements of individual viral particles were measured from micrographs like those shown in panel C. Each circle represents a single virion, and the line denotes the mean with the standard deviation (*n*=35).

### Effect of truncation of the OROV Gc N-terminal variable region on viral infectivity

For the orthobunyavirus, bunyamwera (BUNV), the N-terminal half of Gc is dispensable for viral replication in cell culture (40). Phylogenetic analysis suggests the N-terminus of Gc comprises two variable domains that have novel folds, and recent structural studies reveal that those variable domains correspond to a “head” and “stalk” domain of Gc (16). Deletion of the head domain results in enhanced cell fusion for BUNV, suggesting that this region may act to mask or protect the highly conserved fusion peptide (40). Elimination of the head and stalk domain resulted in a further reduction in viral yield, although the mechanism underlying this is unclear (40). Using the infectious cDNA of VSV-OROV we engineered the analogous variants lacking the “head” domain of Gc (VSV-OROVΔ4) or the “head” and “stalk” domains (VSV-OROVΔ8). Autonomously replicating viruses were recovered from both variants (Figure 2A), although they exhibit a growth defect in cell culture compared to VSV-OROV. Although growth of both truncated variants was attenuated compared to VSV-OROV, VSV-OROVΔ4 reached comparable titers (Figure 2A). Sequence analysis demonstrates that no other changes were present in the GPC gene of VSV-OROV, or VSV-OROVΔ4, but each of the VSV-OROVΔ8 isolates contained additional mutations in Gc (Figure 2B). This result demonstrates that as for BUNV, at least a portion of the N-terminus of Gc is dispensable for infectivity of OROV.

**Figure 2:**
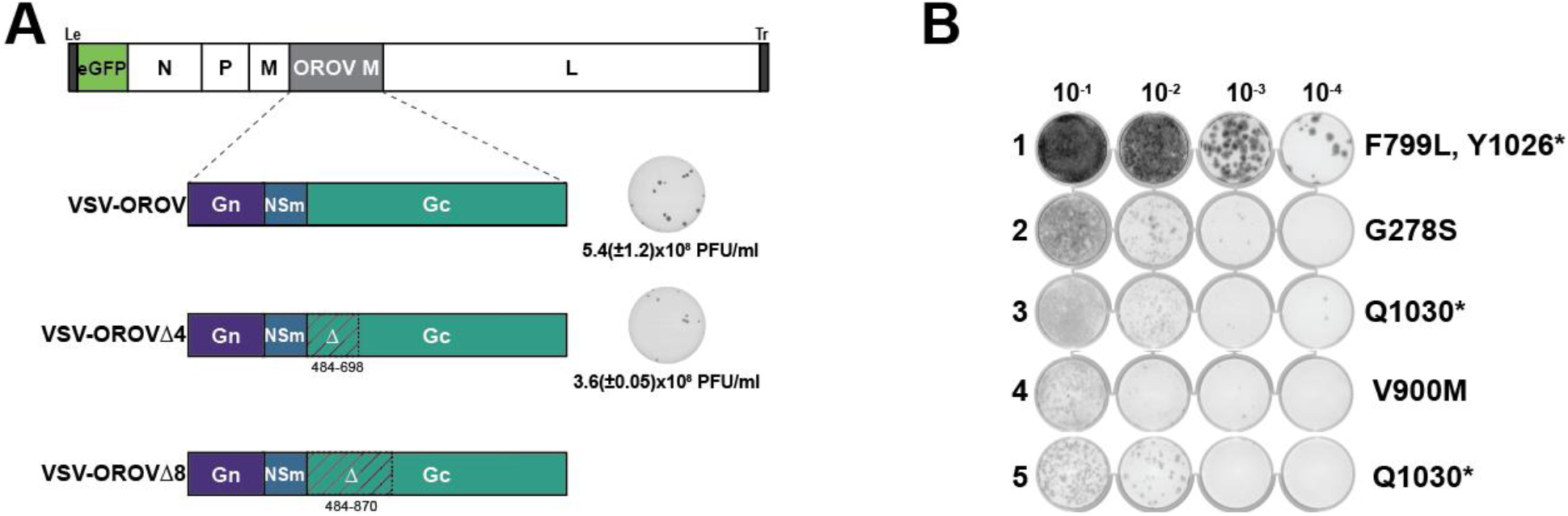
Generation of VSV-OROV chimeras and mutants. (A) Genomic organization of VSV-OROV, VSV-OROVΔ4, and VSV-OROVΔ8. Amino acids 484-698 were deleted from the N-terminal region of Gc to generate VSV-OROVΔ4, and amino acids 484-870 were deleted to generate VSV-OROVΔ8. Plaque assays of VSV-OROV (shown also in Figure 1) and VSV-OROVΔ4 on BSRT7 cells are shown on the right. (B) VSV-OROVΔ8 passaged on BSRT7 cells and the M segment was sequenced by Sanger sequencing. Mutations in the M segment are denoted on the right.

### VSV-OROV chimeras elicit production of neutralizing antibodies in mice

To determine whether the VSV-OROV chimeras were immunogenic, we inoculated BALB/c mice with VSV or the indicated VSV-OROV chimera, boosted the animals 28 days later and examined serum neutralizing titers at days 28 and 35 post inoculation (Figure 3A). Inoculation of mice with VSV-OROV generates sera that specifically neutralize VSV-OROV but not VSV, as measured by cell culture infection assays. Correspondingly, sera from mice inoculated with VSV specifically neutralize VSV but not VSV-OROV, validating that neutralization depends on the surface glycoproteins and not other components of the virion (Figure 3B). All mice produced antibodies capable of neutralizing the relevant virus, whereas sera from sham vaccinated animals failed to neutralize either VSV-OROV or VSV. Mice inoculated with VSV-OROVΔ4 generated sera that neutralized both VSV-OROV and VSV-OROVΔ4 (Figure 3C). The increased sensitivity of VSV-OROVΔ4 to neutralization, suggests loss of the head domain may increase accessibility of important neutralizing epitopes.

**Figure 3:**
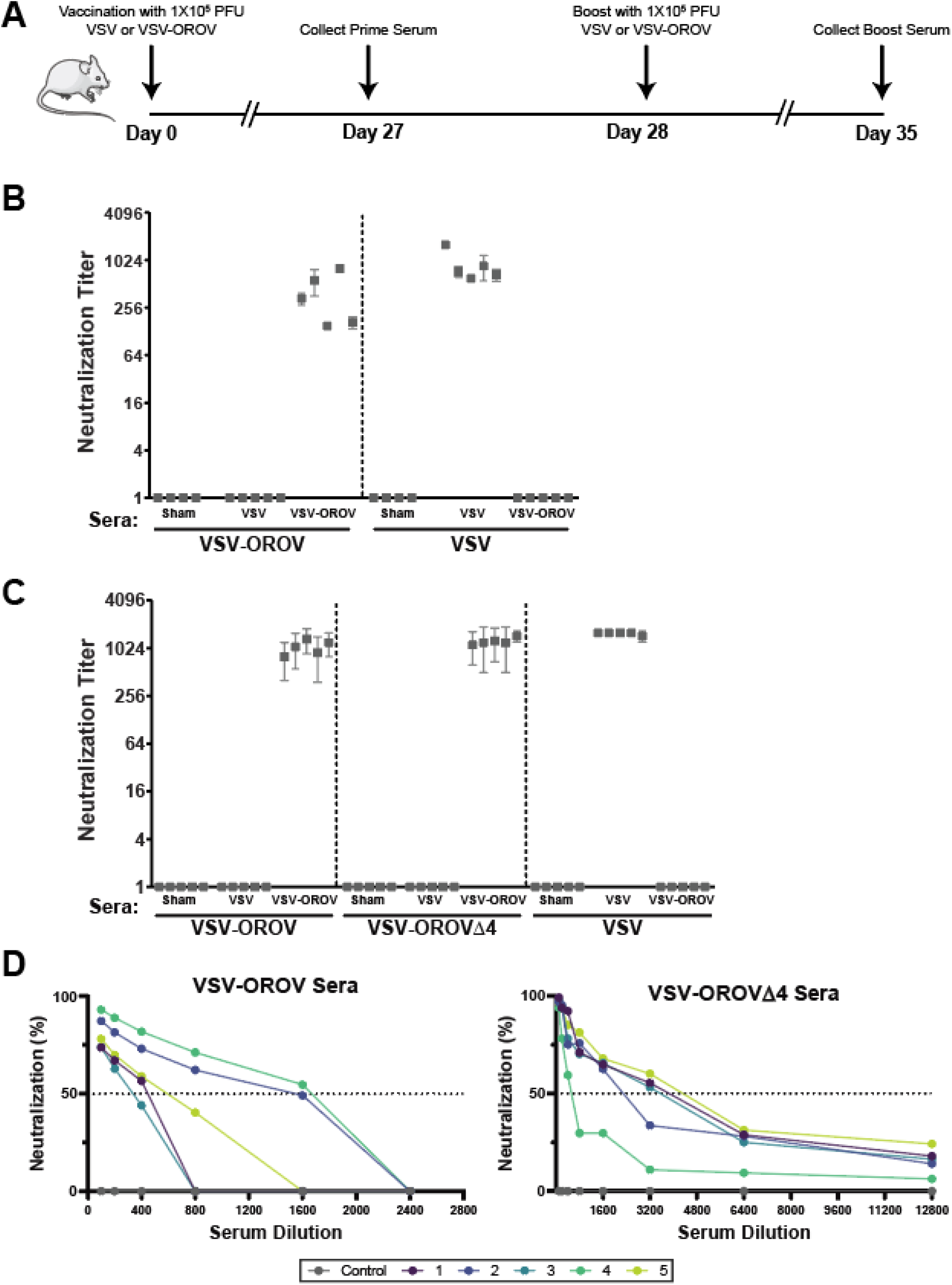
Inoculation of mice with VSV recombinants bearing OROV glycoproteins generates neutralizing antibodies. (A) Inoculation schedule of BALB/c mice. Animals (n= 5 per group) were immunized with VSV-eGFP, VSV-OROV, or VSV-OROVΔ4 and then boosted on day 28. Mice were sacrificed on day 35 and serum was collected. (B) Reciprocals dilution of VSV or VSV-OROV serum that protects cells from 100 TCID50 units of indicated viruses are shown (*n*=3). (C) Reciprocal dilution of VSV or VSV-OROVΔ4 serum that protects cells from viruses indicated (*n*=3). No neutralization at the highest concentration of sera (1:100) was scored as 1. (D) Plaque reduction neutralization assays were performed to assess neutralization of wildtype OROV. Plaques were visually scored after incubation with serial dilutions of mouse serum.

To examine the ability of the mouse sera to neutralize wild-type OROV we used a plaque reduction neutralization titer assay (Figure 3D). Consistent with the ability of the sera to neutralize the respective VSV-chimeras, mice that generated neutralizing antibodies against VSV-OROV neutralized wild-type OROV showing 50% decreases in infectivity at serum dilutions of >1:200-1:600. Consistent with the observed neutralization of the VSV-chimeras, sera from mice inoculated with VSV-OROVΔ4 were more potent than those inoculated with VSV-OROV exhibiting a >50% inhibition of infection at dilutions of 1:400-1:3200 (Figure 3D) We conclude that VSV-OROV results in production of sera that neutralizes wildtype virus and that deletion of the N-terminal portion of Gc further increases neutralizing titers perhaps by targeting additional epitopes.

### VSV-OROV and VSV-OROVΔ4 sera generate antibodies to both Gn and Gc

To identify epitopes recognized by the serum from each immunized mouse we utilized peptide arrays designed to detect antibodies targeting linear epitopes. The tiled array is composed of ~170,000 non-redundant 12-mer peptides from seven arboviruses including OROV (41), and is formatted in a sliding window pattern such that there is an 11 amino acid overlap along the peptide sequence of each virus. The peptides are randomly distributed across the array to prevent location bias. This peptide array was previously used to diagnose Zika virus infection and identified a highly specific Zika epitope within the NS2B protein (41). Profiling of the sera from each mouse inoculated with VSV-OROV identified four peptides in a 150 residue region of the ectodomain of Gc with two of the peptides overlapping by six amino acids (Figure 4A and 4B). This region of Gc corresponds to the N-terminal, “head domain of the variable region of Gc (16) (Figure 4C).

**Figure 4.**
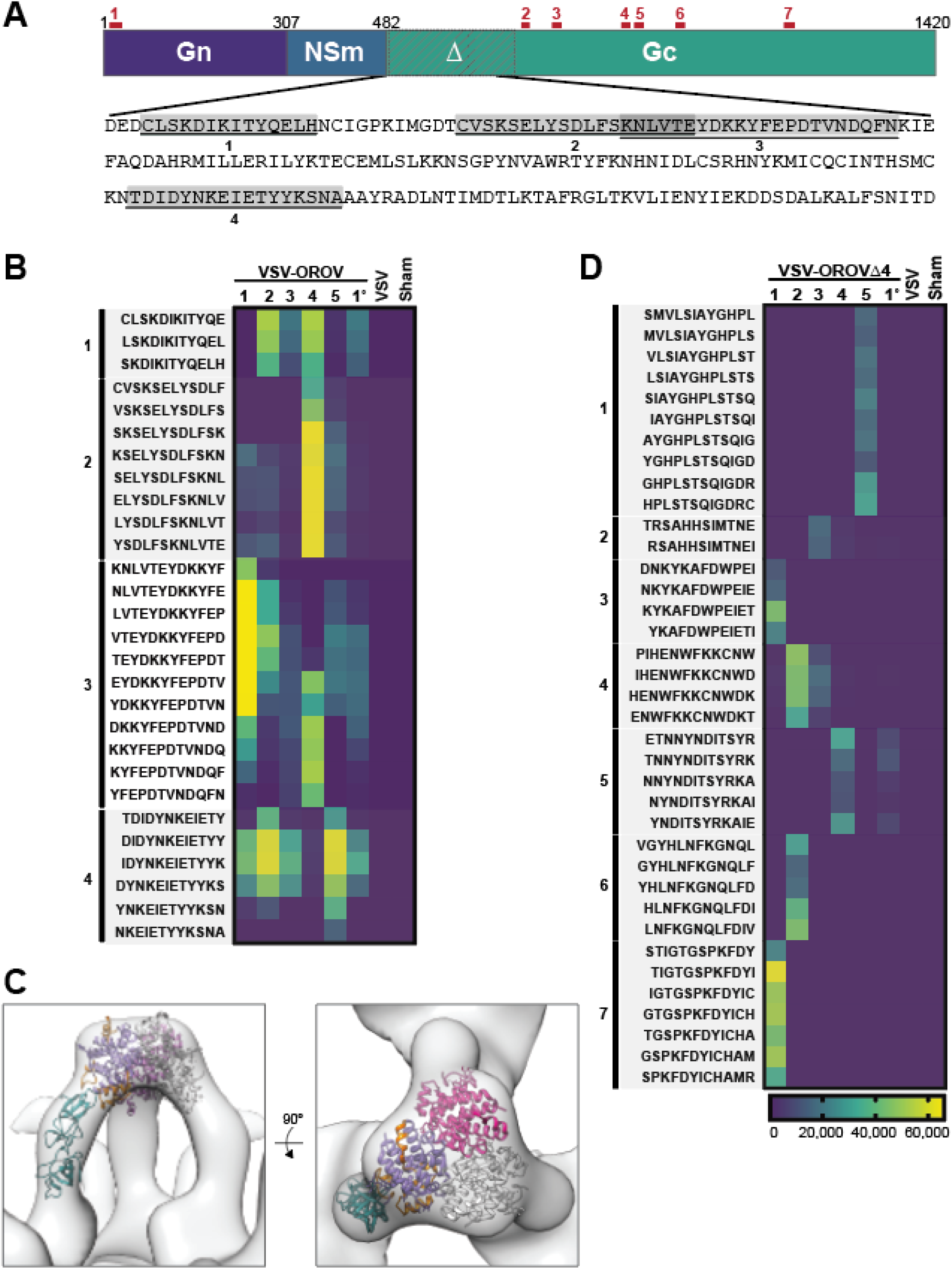
VSV-OROV and VSV-OROVΔ4 antibodies target Gn and Gc. (A) Schematic of OROV M segment showing the two glycoproteins (Gn and Gc) as well as the non-structural, NSm protein. Peptides identified in the peptide array with VSV-OROV are indicated in gray boxes while peptides identified in the VSV-OROVΔ4 peptide array are indicated above the schematic in red lines. (B) Heat map of VSV-OROV peptide array. Analysis on the peptide array was performed with 5 VSV-OROV mouse serum samples, one primary bleed sample, one VSV serum sample, and a sham. Scale is shown in arbitrary units. (C) Mapping of the four identified peptides to the structure of the OROV Gc N-terminal head domain that was determined by Hellert et al. (D) Peptide array from VSV-OROVΔ4 sera, like what is shown in panel B for VSV-OROV.

Mice immunized with VSV-OROVΔ4, lacking the N-terminal region of Gc, generated immune sera that reacted with linear epitopes that target both Gn and Gc of OROV. Specifically, we identified one peptide at the N-terminus of Gn, and six epitopes distributed throughout Gc. Five of the six Gc peptides are located C-terminal to the fusion peptide with 1 peptide mapping 100 residues to the N-terminus of the fusion peptide. Earlier work with Bunyamwera virus defined the C-terminal half of Gc as important for cell fusion and for interaction of Gc and Gn for trafficking through the Golgi (40). When aligned with Gc, two of the five peptides, 974-989 and 1159-1176, map to the C-terminal half of Gc. This data demonstrates that removal of the N-terminal head domain of Gc does not impact the ability of mice to generate neutralizing sera to VSV-OROV and that linear epitopes are generated that target both Gn and Gc.

### Effect of immunization with VSV-OROV chimeras on challenge with wild type OROV

To determine whether VSV-OROV can elicit a protective immune response we employed a 2-dose prime-boost immunization regimen followed by a wild type virus challenge. Briefly, groups of mice (n=5) received intramuscular inoculation of 10^6^ infectious units of VSV, VSV-OROV, or VSV-OROVΔ4, and an equivalent immunization 28 days later. Animals were challenged 7 days later with 10^6^ TCID_50_ of wild type virus and their weights and temperatures examined daily. At 7 days post-challenge, the levels of viral RNA in the brain, liver, and spleen were examined using a quantitative reverse transcription polymerase chain reaction (qRT-PCR) assay (Figure 5A). Mice immunized with VSV or given PBS as a sham lost 5-10% body weight and exhibit a spike in body temperature following challenge. By contrast, animals immunized with VSV-OROV or VSV-OROVΔ4 continued to increase in body weight post-challenge and were not febrile (Figure 5B). We confirmed that mice immunized by this 2-dose regimen of the VSV-OROV chimeras have high neutralizing serum titers against OROV 1 day prior to challenge, whereas the VSV or sham vaccinated animals had no detectable neutralizing titers (Figure 5C).

**Figure 5.**
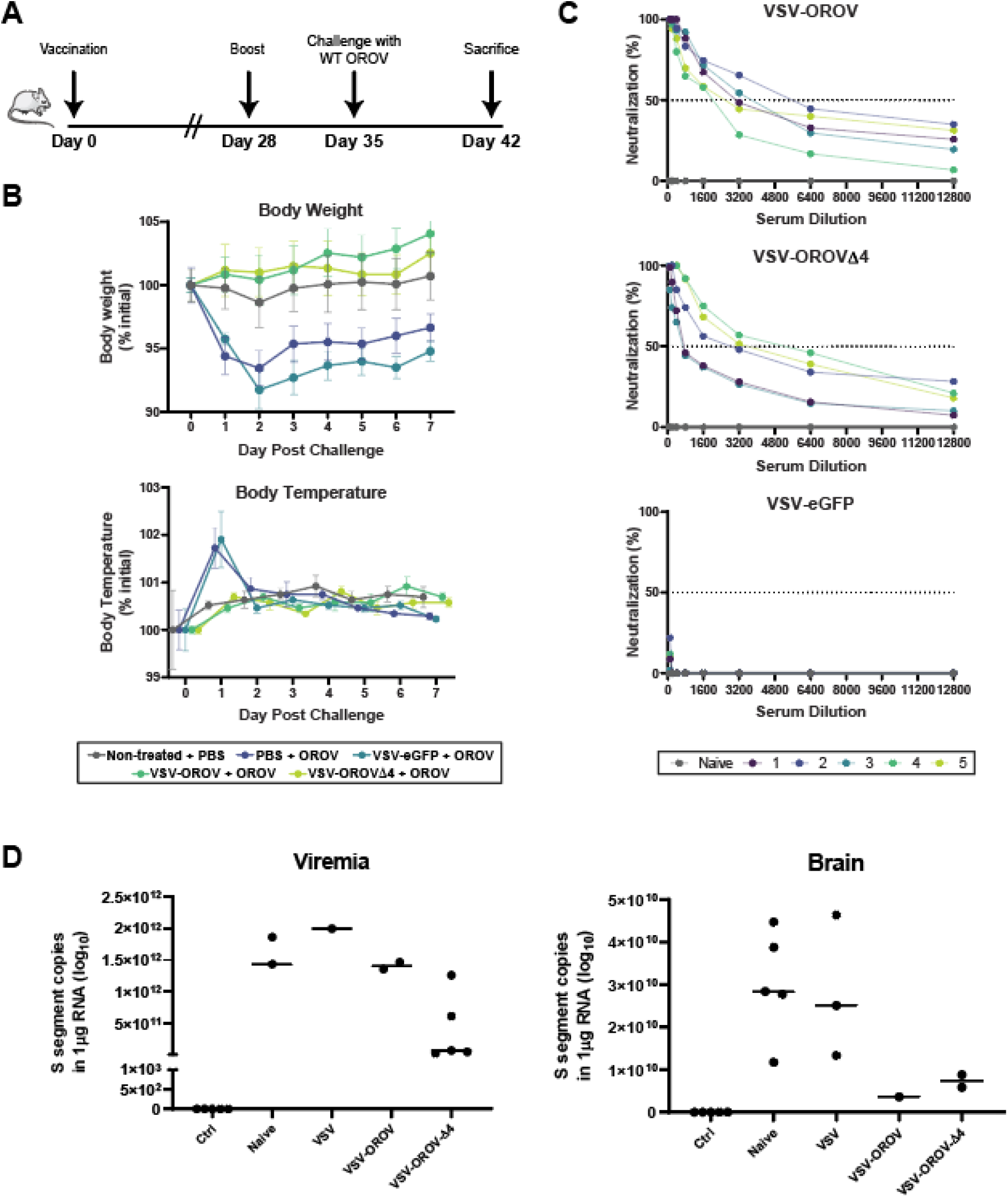
Prime-boost vaccination of mice with VSV recombinants and challenge with OROV. (A) Inoculation schedule of male, 6-week old C57Bl/6 mice. Mice were inoculated on day zero with 10^6^ FFU, boosted on day 28 with the same dose, and then challenged one week later with 10^6^ TCID_50_ OROV (five mice per group). One week after challenge mice were sacrificed. (B) Body weight (top) and temperature (bottom) were assessed each day following challenge with OROV. (C) One day before challenge, VSV-OROV, VSV-OROVΔ4, and VSV sera was collected from mice and tested for neutralization of OROV. (D) Viral loads were assessed by measuring S segment copies in blood and brain samples from mice following sacrifice.

To examine the extent of protection against infection upon challenge we examined viral S segment RNA levels by qRT-PCR 7 days post challenge. We detected S segment RNA copies in the blood of all groups of challenged animals although levels were lowest in the VSV-OROVΔ4 immunized mice. This result demonstrates that the animals develop a viremia following challenge with wildtype virus in the face of a neutralizing serum response (Figure 5D). Analysis of brain tissue demonstrates that mice immunized with VSV-OROV and VSV-OROVΔ4 have reduced viral RNA levels (Figure 5D) consistent with reduced OROV infection.

## Discussion

We report three VSV-chimeric viruses expressing the surface proteins Gn and Gc of the emerging orthobunyavirus, Oropouche, and evaluate those chimeras for immunogenicity and protection against challenge in a mouse model of disease. The VSV chimeras are highly immunogenic in mice, yielding serum neutralizing titers of up to 1:6400 against wild type OROV following a 2-dose intramuscular immunization regimen. Using overlapping peptide arrays we demonstrate that neutralizing serum include antibodies that recognize peptides in the highly variable N-terminus of Gc and demonstrate that elimination of this region results in sera that recognize more conserved regions of Gc. Challenge studies demonstrate that immunized mice are infected although they exhibit reduced clinical symptoms and viral loads. This work demonstrates that neutralizing antisera offer some protection against Oropouche infection and provides new BSL2 tools that can be used to understand the functions of Gn and Gc.

The spread of Oropouche virus in the Amazon region has prompted efforts to advance countermeasures against infection. The inosine monophosphate dehydrogenase inhibitor, ribavirin, has no efficacy against OROV (42), and although innate immune responses control infection of mice by OROV (43), IFN-α treatment offers only limited protection of mice when administered within three hours of infection (42, 44). Earlier efforts to develop vaccines for OROV have included deletion of the NSs and NSm proteins in infectious molecular clones of OROV (3). A similar strategy shows promise for Schmallenberg virus (SBV), an orthobunyavirus that infects ruminants (45). Although correlates of immunity are not completely understood for many bunyaviruses, vaccines targeting the glycoproteins appear to illicit strong immune responses for at least some bunyaviruses. DNA vaccines expressing Gn and Gc protected mice from challenge with RVFV, while Gn and Gc of CCHFV were not sufficient to confer protection (46). Subunit vaccines comprising the head domain of the orthobunyavirus SBV Gc protected mice from clinical symptoms of disease, while a misfolded version of the head domain did not (47).

Here we demonstrate that immunization of mice with VSV-OROV chimeras generates a strong neutralization serum response against wild type OROV. That neutralizing serum response correlates with some protection against infection as evidenced by reduced clinical signs – such as weight loss and a febrile response – although viral infection was not prevented. Analysis of the serum neutralizing antibody response using linear peptide arrays identified four prominent epitopes within the N-terminal region of Gc. All four of those peptides lie within the “head” domain of the Gc N-terminal region (16). Recognition of the OROV head domain fits with earlier work on La Crosse virus (LACV) and SBV have also shown that a large proportion of glycoprotein specific antibodies are targeted to the Gc head domain (20, 21). Immunization of mice with the purified head, or head and stalk domains of Gc protects against infection with SBV (16, 47), demonstrating that for some orthobunyaviruses targeting this region of Gc may be an effective strategy for vaccine development. Although we did not directly test whether the N terminus of Gc of OROV can elicit the induction of protective immune responses alone, its loss did not diminish neutralizing serum titers and instead led to the identification of additional epitopes within Gc. Though we observe a modest distinction in the extent of infection upon challenge of mice immunized with VSV-OROV and VSV-OROVΔ4, we do not understand the basis for this distinction. Development of Oropouche vaccines will require a deeper understanding of correlates of immune protection, and whether immune protection is achieved following natural infection. The VSV-OROV chimeras described here may aid in this analysis by providing useful BSL2 tools to study Gn and Gc function and their inhibition.

## Materials and Methods

### Cell Lines

African green monkey Vero cells and BSRT7 Syrian golden hamster kidney cells (generously provided by K. Conzelmann (48) were maintained at 37°C and 5% CO_2_ in Dulbecco’s modified Eagle medium (DMEM) supplemented with 10% fetal bovine serum (FBS).

### VSV Recombinant Generation, Growth, Purification, and Titration

A codon-optimized version of the M segment of OROV BeAn19991 sequence was ordered from GenScript (Piscataway, NJ). The OROV M segment was cloned into pVSV-ΔG-eGFP (39) using MluI and NotI sites. To generate OROVΔ4, site directed mutagenesis was performed using primers GATATCAACCTGGGCAGG and CTCGTCGGCGTACACTGT. VSV-OROV and VSV-OROVΔ4 were generated by transfecting the genomic plasmid along with plasmids containing VSV N, P, L, and G into BSRT7 cells, as previously described (23). VSV containing eGFP (enhanced green fluorescent protein) was previously generated (39). All viruses were grown on BSRT7 cells in DMEM containing 2% FBS and penicillin-streptomycin/kanamycin. Viral supernatants were titered by plaque assay as previously described (49) or gradient purified on a 15-45% sucrose gradient and then tittered. GFP expressing plaques were visualized on a Typhoon 9400 imager. Purified virus was run on a 10% (wt/vol) acrylamide 0.13% (wt/vol) bis-acrylamide gel. Gels were stained with Coomassie reagent.

### Electron Microscopy

VSV and VSV-OROV were deposited onto carbon-coated copper grids and then stained with 2% (wt/vol) phosphotungstic acid (PTA) in H_2_O (pH 7.5). A Technai G^2^ Spirit BioTwin transmission electron microscope (FEI, Hillsboro, OR) was used to visualize particles. Length and width of viral particles was measured using ImageJ.

### Generation of Neutralizing Sera

Six-week old male BALB/c mice (Taconic Farms) were injected intramuscularly (IM) with 10^5^ PFU of VSV, VSV-OROV, VSV-OROVΔ4 or sham on day zero. On day 27, tail vein blood samples were collected and on day 28 mice were boosted with an additional dose of 10^5^ PFU of virus. Seven days later on day 35 mice were sacrificed and blood samples were collected by cardiac puncture. Serum was separated by centrifugation at 1500xg for 10 minutes at room temperature and then heated at 56°C for 30’ to inactivate complement.

### Serum Neutralization Assays

Heat inactivated serum samples from mice were diluted in DMEM in 2-fold dilution series ranging from 1:100 to 1:1600 in duplicate on a 96-well plate. 100 TCID50 units of VSV, VSV-OROV, or VSV-OROVΔ4 were incubated with each serum dilution in well for 1.5hr at 37°C. Following incubation of virus and serum, 30,000 BSRT7 cells were added to each well and incubated at 34°C for 48 hours. 96-well plates were scanned on a Typhoon FLA9500 Imager and dilutions lacking any GFP signal was recorded. Neutralization assays were performed three times.

### OROV Sera Peptide Arrays

VSV, VSV-OROV, VSV-OROVΔ4, and sham sera samples were analyzed using a previously described arboviral peptide array (41, 50).

### WT OROV Stock Production

OROV strain BeAn19991 was originally donated from Luiz Tadeu Figueiredo and propagated by serial passages in Vero cells by routine methods using DMEM. Virus titration was performed by plaque assay in Vero cells plated at 4 x10^4^ cells/well in 48-well plates one day prior to infection. After 1 h incubation with virus, cells were replenished with DMEM supplemented with 2% v/v FBS, 1% v/v antibiotics and 1% v/v carboxymethyl cellulose (CMC) (Sigma-Aldrich), and incubated at 37°C and 5% CO_2_ for 4 days. Cells were fixed with 4% formaldehyde for 20 min at room temperature, washed in Phosphate Buffered Saline (PBS) (Gibco) and stained with 20% v/v ethanol-violet crystal solution for 15 min.

### OROV PRNT_50_

The titer of neutralizing antibodies was determined on serum obtained on day 28 after immunization with VSV, VSV-OROV or VSV-OROVΔ4 mice by a standard plaque reduction neutralization assay. Plaques were scored visually after incubation with serial dilutions of the mouse serum, and the 50% plaque reduction neutralization titer (PRNT_50_) was determined (51).

### Immunization and OROV Challenge

A total of 25 male C57Bl/6 6-week-old mice were assigned to 5 groups: two sham groups were administered sterile PBS 1x, and three groups were immunized with VSV, VSV-OROV or VSV-OROVΔ4. Injections were made subcutaneously in the dorsal lumbar area with 10^6^ FFU in a volume of 50uL. At 28 days post-vaccination a booster immunization was administered to animals with the same virus dose or volume of PBS. One week later the VSV recombinant-immunized and one sham-inoculated group were challenged with 10^6^ TCID_50_ OROV wild type BeAn19991 subcutaneously. The other sham-inoculated group was mock-injected. Mice were followed for weight loss and temperature variation once per day for seven days following challenge. All injections with virus were performed under anesthesia of 1.5% isoflurane.

### WT OROV Serum and organ sampling

Serum from animals was collected by retrieving blood from the caudal vein. Total blood was centrifuged at 5.000 rpm for 5 min and clear serum was collected. Seven days after challenge, mice were sacrificed and perfused through the ascending aorta with sterile PBS (pH 7.4). Brain, liver, spleen, kidneys, and spinal cord were obtained by dissection under aseptic conditions. Organs were sections, weighed, and stored in TRIzol (Thermo Scientific) at −70°C. One sample of each brain was also collected for immunohistochemistry.

### RT qPCR

Tissues were dissected, weighed, and homogenized in sterile PBS using a TissueLyser II (Qiagen). For viral RNA quantification, samples were extracted using a QIAamp Viral RNA Mini Kit (Qiagen) according to the manufacturer’s recommendations. Reverse transcription was performed using random primers and the High Capacity Reverse Transcriptase kit (Thermo Fisher), according to manufacturer’s instructions. Primers were designed to detect a 100 bp region of OROV nucleocapsid from the small segment (S-OROV-rev: 5’-TTGCGTCACCATCATTCCAA-3’; S-OROV-fow: 5’-TACCCAGATGCGATCACCAA-3’), using Sybr Green (Kappa Biosystems). The reaction was performed in the StepOnePlus PCR System (Applied Biosystems). Standard curves were generated using a plasmid containing the entire OROV small segment antigenome (pTVT-S described in Tilston-Lunel, 2015). Each sample was assayed in duplicate and plotted the mean value (3).

### Animal Use and Ethical Statement

All VSV recombinant animal experiments and housing were conducted in accordance with protocols approved by the Harvard Medical Area Standing Committee on Animals. All WT OROV studies were approved by the University of São Paulo Committee on Care and Use of Laboratory Animals (protocol #194/2019). 6-week-old C57Bl/6 mice were obtained from the Central Animal Facility of the University of São Paulo, School of Medicine, in Ribeirão Preto, SP, Brazil. Infected animals were maintained in the Virology Research Center – FMRP USP animal facility. All animals were kept in accordance with guidelines of the University of São Paulo Committee on Care and Use of Laboratory Animals.

## Acknowledgements

This work was supported by NIH grants U19 AI109740, U19 AI109740, AI057552, and AI098681 as well as FAPESP grant 2016/06490-8 and funds from the Coordination for the Improvement of Higher Education Personnel.

## References

1. Sakkas H, Bozidis P, Franks A, Papadopoulou C. 2018. Oropouche Fever: A Review. Viruses 10.

2. Romero-Alvarez D, Escobar LE. 2018. Oropouche fever, an emergent disease from the Americas. Microbes Infect 20:135–146.

3. Tilston-Lunel NL, Acrani GO, Randall RE, Elliott RM. 2015. Generation of Recombinant Oropouche Viruses Lacking the Nonstructural Protein NSm or NSs. J Virol 90:2616–27.

4. Figueiredo LT. 2007. Emergent arboviruses in Brazil. Rev Soc Bras Med Trop 40:224–9.

5. Forshey BM, Guevara C, Laguna-Torres VA, Cespedes M, Vargas J, Gianella A, Vallejo E, Madrid C, Aguayo N, Gotuzzo E, Suarez V, Morales AM, Beingolea L, Reyes N, Perez J, Negrete M, Rocha C, Morrison AC, Russell KL, Blair PJ, Olson JG, Kochel TJ. 2010. Arboviral etiologies of acute febrile illnesses in Western South America, 2000-2007. PLoS Negl Trop Dis 4:e787.

6. Navarro JC, Giambalvo D, Hernandez R, Auguste AJ, Tesh RB, Weaver SC, Montañez H, Liria J, Lima A, Travassos da Rosa JF, da Silva SP, Vasconcelos JM, Oliveira R, Vianez JL, Jr., Nunes MR. 2016. Isolation of Madre de Dios Virus (Orthobunyavirus; Bunyaviridae), an Oropouche Virus Species Reassortant, from a Monkey in Venezuela. Am J Trop Med Hyg 95:328–38.

7. Manock SR, Jacobsen KH, de Bravo NB, Russell KL, Negrete M, Olson JG, Sanchez JL, Blair PJ, Smalligan RD, Quist BK, Espín JF, Espinoza WR, MacCormick F, Fleming LC, Kochel T. 2009. Etiology of acute undifferentiated febrile illness in the Amazon basin of Ecuador. Am J Trop Med Hyg 81:146–51.

8. Aguilar PV, Barrett AD, Saeed MF, Watts DM, Russell K, Guevara C, Ampuero JS, Suarez L, Cespedes M, Montgomery JM, Halsey ES, Kochel TJ. 2011. Iquitos virus: a novel reassortant Orthobunyavirus associated with human illness in Peru. PLoS Negl Trop Dis 5:e1315.

9. Baisley KJ, Watts DM, Munstermann LE, Wilson ML. 1998. Epidemiology of endemic Oropouche virus transmission in upper Amazonian Peru. Am J Trop Med Hyg 59:710–6.

10. Pinheiro FP, Travassos da Rosa AP, Travassos da Rosa JF, Ishak R, Freitas RB, Gomes ML, LeDuc JW, Oliva OF. 1981. Oropouche virus. I. A review of clinical, epidemiological, and ecological findings. Am J Trop Med Hyg 30:149–60.

11. Travassos da Rosa JF, de Souza WM, Pinheiro FP, Figueiredo ML, Cardoso JF, Acrani GO, Nunes MRT. 2017. Oropouche Virus: Clinical, Epidemiological, and Molecular Aspects of a Neglected Orthobunyavirus. Am J Trop Med Hyg 96:1019–1030.

12. Pinheiro FP, Travassos da Rosa AP, Gomes ML, LeDuc JW, Hoch AL. 1982. Transmission of Oropouche virus from man to hamster by the midge Culicoides paraensis. Science 215:1251–3.

13. Romero-Alvarez D, Escobar LE. 2017. Vegetation loss and the 2016 Oropouche fever outbreak in Peru. Mem Inst Oswaldo Cruz 112:292–298.

14. Elliott RM, Blakqori G. 2011. Molecular biology of orthobunyaviruses. Caister Academic Press, Norfolk, U.K.

15. Garry CE, Garry RF. 2004. Proteomics computational analyses suggest that the carboxyl terminal glycoproteins of Bunyaviruses are class II viral fusion protein (beta-penetrenes). Theor Biol Med Model 1:10.

16. Hellert J, Aebischer A, Wernike K, Haouz A, Brocchi E, Reiche S, Guardado-Calvo P, Beer M, Rey FA. 2019. Orthobunyavirus spike architecture and recognition by neutralizing antibodies. Nat Commun 10:879.

17. Grady LJ, Kinch W. 1985. Two monoclonal antibodies against La Crosse virus show host-dependent neutralizing activity. J Gen Virol 66 (Pt 12):2773–6.

18. Ludwig GV, Israel BA, Christensen BM, Yuill TM, Schultz KT. 1991. Monoclonal antibodies directed against the envelope glycoproteins of La Crosse virus. Microb Pathog 11:411–21.

19. Schuh T, Schultz J, Moelling K, Pavlovic J. 1999. DNA-based vaccine against La Crosse virus: protective immune response mediated by neutralizing antibodies and CD4+ T cells. Hum Gene Ther 10:1649–58.

20. Kingsford L, Ishizawa LD, Hill DW. 1983. Biological activities of monoclonal antibodies reactive with antigenic sites mapped on the G1 glycoprotein of La Crosse virus. Virology 129:443–55.

21. Roman-Sosa G, Brocchi E, Schirrmeier H, Wernike K, Schelp C, Beer M. 2016. Analysis of the humoral immune response against the envelope glycoprotein Gc of Schmallenberg virus reveals a domain located at the amino terminus targeted by mAbs with neutralizing activity. J Gen Virol 97:571–580.

22. Fields BN, Knipe DM, Howley PM. 2013. Fields virology, 6th ed. Wolters Kluwer Health/Lippincott Williams & Wilkins, Philadelphia.

23. Whelan SP, Ball LA, Barr JN, Wertz GT. 1995. Efficient recovery of infectious vesicular stomatitis virus entirely from cDNA clones. Proc Natl Acad Sci U S A 92:8388–92.

24. van den Pol AN, Mao G, Chattopadhyay A, Rose JK, Davis JN. 2017. Chikungunya, Influenza, Nipah, and Semliki Forest Chimeric Viruses with Vesicular Stomatitis Virus: Actions in the Brain. J Virol 91.

25. Geisbert TW, Jones S, Fritz EA, Shurtleff AC, Geisbert JB, Liebscher R, Grolla A, Ströher U, Fernando L, Daddario KM, Guttieri MC, Mothé BR, Larsen T, Hensley LE, Jahrling PB, Feldmann H. 2005. Development of a new vaccine for the prevention of Lassa fever. PLoS Med 2:e183.

26. Garbutt M, Liebscher R, Wahl-Jensen V, Jones S, Möller P, Wagner R, Volchkov V, Klenk HD, Feldmann H, Ströher U. 2004. Properties of replication-competent vesicular stomatitis virus vectors expressing glycoproteins of filoviruses and arenaviruses. J Virol 78:5458–65.

27. Brown KS, Safronetz D, Marzi A, Ebihara H, Feldmann H. 2011. Vesicular stomatitis virus-based vaccine protects hamsters against lethal challenge with Andes virus. J Virol 85:12781–91.

28. Roberts A, Kretzschmar E, Perkins AS, Forman J, Price R, Buonocore L, Kawaoka Y, Rose JK. 1998. Vaccination with a recombinant vesicular stomatitis virus expressing an influenza virus hemagglutinin provides complete protection from influenza virus challenge. J Virol 72:4704–11.

29. Henao-Restrepo AM, Longini IM, Egger M, Dean NE, Edmunds WJ, Camacho A, Carroll MW, Doumbia M, Draguez B, Duraffour S, Enwere G, Grais R, Gunther S, Hossmann S, Kondé MK, Kone S, Kuisma E, Levine MM, Mandal S, Norheim G, Riveros X, Soumah A, Trelle S, Vicari AS, Watson CH, Kéïta S, Kieny MP, Røttingen JA. 2015. Efficacy and effectiveness of an rVSV-vectored vaccine expressing Ebola surface glycoprotein: interim results from the Guinea ring vaccination cluster-randomised trial. Lancet 386:857–66.

30. Henao-Restrepo AM, Camacho A, Longini IM, Watson CH, Edmunds WJ, Egger M, Carroll MW, Dean NE, Diatta I, Doumbia M, Draguez B, Duraffour S, Enwere G, Grais R, Gunther S, Gsell PS, Hossmann S, Watle SV, Kondé MK, Kéïta S, Kone S, Kuisma E, Levine MM, Mandal S, Mauget T, Norheim G, Riveros X, Soumah A, Trelle S, Vicari AS, Røttingen JA, Kieny MP. 2017. Efficacy and effectiveness of an rVSV-vectored vaccine in preventing Ebola virus disease: final results from the Guinea ring vaccination, open-label, cluster-randomised trial (Ebola Ça Suffit!). Lancet 389:505–518.

31. Huttner A, Dayer JA, Yerly S, Combescure C, Auderset F, Desmeules J, Eickmann M, Finckh A, Goncalves AR, Hooper JW, Kaya G, Krähling V, Kwilas S, Lemaître B, Matthey A, Silvera P, Becker S, Fast PE, Moorthy V, Kieny MP, Kaiser L, Siegrist CA. 2015. The effect of dose on the safety and immunogenicity of the VSV Ebola candidate vaccine: a randomised double-blind, placebo-controlled phase 1/2 trial. Lancet Infect Dis 15:1156–1166.

32. Furuyama W, Reynolds P, Haddock E, Meade-White K, Quynh Le M, Kawaoka Y, Feldmann H, Marzi A. 2020. A single dose of a vesicular stomatitis virus-based influenza vaccine confers rapid protection against H5 viruses from different clades. NPJ Vaccines 5:4.

33. Case JB, Rothlauf PW, Chen RE, Kafai NM, Fox JM, Smith BK, Shrihari S, McCune BT, Harvey IB, Keeler SP, Bloyet LM, Zhao H, Ma M, Adams LJ, Winkler ES, Holtzman MJ, Fremont DH, Whelan SPJ, Diamond MS. 2020. Replication-Competent Vesicular Stomatitis Virus Vaccine Vector Protects against SARS-CoV-2-Mediated Pathogenesis in Mice. Cell Host Microbe 28:465–474.e4.

34. Carette JE, Raaben M, Wong AC, Herbert AS, Obernosterer G, Mulherkar N, Kuehne AI, Kranzusch PJ, Griffin AM, Ruthel G, Dal Cin P, Dye JM, Whelan SP, Chandran K, Brummelkamp TR. 2011. Ebola virus entry requires the cholesterol transporter Niemann-Pick C1. Nature 477:340–3.

35. Piccinotti S, Whelan SP. 2016. Rabies Internalizes into Primary Peripheral Neurons via Clathrin Coated Pits and Requires Fusion at the Cell Body. PLoS Pathog 12:e1005753.

36. Robinson-McCarthy LR, McCarthy KR, Raaben M, Piccinotti S, Nieuwenhuis J, Stubbs SH, Bakkers MJG, Whelan SPJ. 2018. Reconstruction of the cell entry pathway of an extinct virus. PLoS Pathog 14:e1007123.

37. Jae LT, Raaben M, Herbert AS, Kuehne AI, Wirchnianski AS, Soh TK, Stubbs SH, Janssen H, Damme M, Saftig P, Whelan SP, Dye JM, Brummelkamp TR. 2014. Virus entry. Lassa virus entry requires a trigger-induced receptor switch. Science 344:1506–10.

38. Jones SM, Feldmann H, Ströher U, Geisbert JB, Fernando L, Grolla A, Klenk HD, Sullivan NJ, Volchkov VE, Fritz EA, Daddario KM, Hensley LE, Jahrling PB, Geisbert TW. 2005. Live attenuated recombinant vaccine protects nonhuman primates against Ebola and Marburg viruses. Nat Med 11:786–90.

39. Wong AC, Sandesara RG, Mulherkar N, Whelan SP, Chandran K. 2010. A forward genetic strategy reveals destabilizing mutations in the Ebolavirus glycoprotein that alter its protease dependence during cell entry. J Virol 84:163–75.

40. Shi X, Goli J, Clark G, Brauburger K, Elliott RM. 2009. Functional analysis of the Bunyamwera orthobunyavirus Gc glycoprotein. J Gen Virol 90:2483–2492.

41. Mishra N, Caciula A, Price A, Thakkar R, Ng J, Chauhan LV, Jain K, Che X, Espinosa DA, Montoya Cruz M, Balmaseda A, Sullivan EH, Patel JJ, Jarman RG, Rakeman JL, Egan CT, Reusken C, Koopmans MPG, Harris E, Tokarz R, Briese T, Lipkin WI. 2018. Diagnosis of Zika Virus Infection by Peptide Array and Enzyme-Linked Immunosorbent Assay. MBio 9.

42. Livonesi MC, De Sousa RL, Badra SJ, Figueiredo LT. 2006. In vitro and in vivo studies of ribavirin action on Brazilian Orthobunyavirus. Am J Trop Med Hyg 75:1011–6.

43. Proenca-Modena JL, Sesti-Costa R, Pinto AK, Richner JM, Lazear HM, Lucas T, Hyde JL, Diamond MS. 2015. Oropouche virus infection and pathogenesis are restricted by MAVS, IRF-3, IRF-7, and type I interferon signaling pathways in nonmyeloid cells. J Virol 89:4720–37.

44. Livonesi MC, de Sousa RL, Badra SJ, Figueiredo LT. 2007. In vitro and in vivo studies of the Interferon-alpha action on distinct Orthobunyavirus. Antiviral Res 75:121–8.

45. Kraatz F, Wernike K, Hechinger S, König P, Granzow H, Reimann I, Beer M. 2015. Deletion mutants of Schmallenberg virus are avirulent and protect from virus challenge. J Virol 89:1825–37.

46. Spik K, Shurtleff A, McElroy AK, Guttieri MC, Hooper JW, SchmalJohn C. 2006. Immunogenicity of combination DNA vaccines for Rift Valley fever virus, tick-borne encephalitis virus, Hantaan virus, and Crimean Congo hemorrhagic fever virus. Vaccine 24:4657–66.

47. Wernike K, Aebischer A, Roman-Sosa G, Beer M. 2017. The N-terminal domain of Schmallenberg virus envelope protein Gc is highly immunogenic and can provide protection from infection. Sci Rep 7:42500.

48. Buchholz UJ, Finke S, Conzelmann KK. 1999. Generation of bovine respiratory syncytial virus (BRSV) from cDNA: BRSV NS2 is not essential for virus replication in tissue culture, and the human RSV leader region acts as a functional BRSV genome promoter. J Virol 73:251–9.

49. Cureton DK, Massol RH, Saffarian S, Kirchhausen TL, Whelan SP. 2009. Vesicular stomatitis virus enters cells through vesicles incompletely coated with clathrin that depend upon actin for internalization. PLoS Pathog 5:e1000394.

50. Tokarz R, Mishra N, Tagliafierro T, Sameroff S, Caciula A, Chauhan L, Patel J, Sullivan E, Gucwa A, Fallon B, Golightly M, Molins C, Schriefer M, Marques A, Briese T, Lipkin WI. 2018. A multiplex serologic platform for diagnosis of tick-borne diseases. Sci Rep 8:3158.

51. Proenca-Modena JL, Hyde JL, Sesti-Costa R, Lucas T, Pinto AK, Richner JM, Gorman MJ, Lazear HM, Diamond MS. 2016. Interferon-Regulatory Factor 5-Dependent Signaling Restricts Orthobunyavirus Dissemination to the Central Nervous System. J Virol 90:189–205.

